# Identification and Structural Characterization of a mutant KRAS-G12V specific TCR restricted by HLA-A3

**DOI:** 10.1101/2024.02.01.578367

**Authors:** Malcolm J. W. Sim, Ken-ichi Hanada, Zachary Stotz, Zhiya Yu, Jinghua Lu, Paul Brennan, Max Quastel, Geraldine M. Gillespie, Eric O. Long, James C. Yang, Peter D. Sun

## Abstract

Mutations in KRAS are some of the most common across multiple cancer types and are thus attractive targets for therapy. Recent studies demonstrated that mutant KRAS generates immunogenic neoantigens that can be targeted in adoptive T cell therapy in metastatic diseases. To expand mutant KRAS specific immunotherapies, it is critical to identify additional HLA-I allotypes that can present KRAS neoantigens and their cognate T cell receptors (TCR). Here, we identified a murine TCR specific to a KRAS-G12V neoantigen (^7^VVVGAVGVGK^16^) using a vaccination approach with transgenic mice expressing the common HLA-I allotype, HLA-A*03:01 (HLA-A3). This TCR demonstrated exquisite specificity for mutant G12V and not Wt KRAS peptides. To investigate the molecular basis for neoantigen recognition by This TCR, we determined its structure in complex with HLA-A3(G12V). G12V-TCR CDR3β and CDR1 β formed a hydrophobic pocket to interact with p6 Val of the G12V but not Wt KRAS peptide. To improve the tumor sensitivity of This TCR, we designed rational substitutions to improve TCR:HLA-A3 contacts. Two substitutions exhibited modest improvements in TCR binding to HLA-A3 (G12V), but did not sufficiently improve T cell sensitivity for further clinical development. Our study provides mechanistic insight into how TCRs detect neoantigens and reveals the challenges in targeting KRAS-G12V mutations. [203]

## Introduction

Adoptive T cell therapy treats cancer by the transfer of tumour specific T cells [1]. An attractive strategy is to target ‘public neoantigens’, which derive from common genetic abnormalities found in multiple cancers and patients such as mutations in p53 and KRAS [2–4]. Encouraging this approach are two demonstrations that adoptive transfer of T cells specific for the common G12D (Gly to Asp at position 12) mutation in KRAS led to tumour regression in patients with metastatic disease [5, 6]. In both cases, the patient carried the HLA class I allotype HLA-C*08:02, which is essential for presentation of the two KRAS-G12D neoantigens targeted by the therapeutic T cell receptors (TCR)[7]. Limiting the expansion of this approach is the population frequency of *HLA-C*08:02*, which is approximately 5-10% worldwide, restricting the number of eligible patients to approximately 4,000 per year (in the US), despite the high frequency of KRAS-G12D mutations in certain cancers [3, 4]. Therefore, to expand these therapies, it is imperative to identify mutant KRAS specific TCRs restricted by additional HLA-I and HLA-II allotypes. More generally, it is also necessary to understand the molecular basis for how TCRs detect neoantigens, given that they differ from wild type (WT) ‘self’ sequences by only single amino acid substitutions [2, 8]. Furthermore, it is unclear if there are any molecular or biophysical features of neoantigen specific TCRs that correlate with *in vitro* tumour reactivity or *in vivo* clinical efficacy [2].

Mutant KRAS specific TCRs have been identified from patient derived tumor infiltrating lymphocytes (TILs) cultures and PBMCs from healthy individuals [9–14]. Another strategy is to isolate neoantigen specific TCRs from HLA transgenic mice vaccinated against specific neoantigens [4, 15]. Taking this approach, we identified an HLA-A*11:01 (HLA-A11) restricted TCR specific for a G12V-KRAS neoantigen, demonstrating reactivity to multiple tumour lines carrying G12V-KRAS and *in vivo* efficacy against an established tumour in an adoptive transfer model [15]. HLA-A*03:01 (HLA-A3) and HLA-A11 are highly related, sharing 98% (334/341) protein sequence identity and present similar peptides utilizing the same positively charged C-terminal anchor residue [16–18]. Further, recent evidence showed that HLA-A3 can also present G12V-KRAS neoantigens and be targeted by T cells and peptide-specific mAbs [11, 19, 20]. Highlighting the importance of identifying reagents that target this mutation, the incidence of cancers carrying the G12V-KRAS mutation in HLA-A3+ individuals is approximately 10,000 per year (in the US) [3]. In the present study, we identified a G12V-KRAS specific TCR from vaccinated HLA-A3 transgenic mice and determined the structural basis for G12V-KRAS recognition by this HLA-A3 restricted TCR.

## Results

### Identification of KRAS-G12V specific HLA-A3 restricted TCR from HLA-A3 transgenic mice by vaccination

Much like HLA-A11, HLA-A3 is predicted to bind G12V-KRAS neoantigens (Table 1). To identify HLA-A3 restricted G12V-KRAS specific TCRs we took a similar approach to our successful identification of HLA-A11 restricted G12V-KRAS specific TCRs using HLA transgenic mice [15]. HLA-A3 transgenic mice (referred to as A3-Kb mice) were generated as previously described [15]. Mice were initially immunized with adenovirus encoding KRAS-G12V_1-30_ followed by three mRNA vaccines encoding MVVVGA**V**GVGK. Subsequently, splenocytes from vaccinated animals were cultured *in vitro* in 24-well plates with either the G12V-9mer (^8^VVGA**V**GVGK^16^) or G12V-10mer (^7^VVVGA**V**GVGK^16^) peptide. Following this, two additional rounds of restimulation were conducted, first with ConA-treated lymphoblasts from A3-Kb mice loaded with the cognate peptide, while the second round utilized EL-4 cells transduced with A3-Kb and KRAS-G12V_1-30_ (EL4-A3-G12V) (Fig. 1A). In a subsequent assay measuring IFN-γ secretion, cells in two wells, B5 and C6, exhibited reactivity to EL4-A3 cells loaded with the G12V-10mer peptide and EL4-A3-G12V (Fig. 1B). Notably, they exhibited no reactivity against G12V 9-mer peptide or EL4-A3-G12D. Further analysis through 5’ RACE and TCR amplification by PCR revealed that the most dominant clone in these wells shared the same TRAV12-3*03 with CDR3α CALSEGGNYKYVF and TRBV1*01 with CDR3 CTCSARHSAETLYF (Fig. 1C).

**Figure 1.**
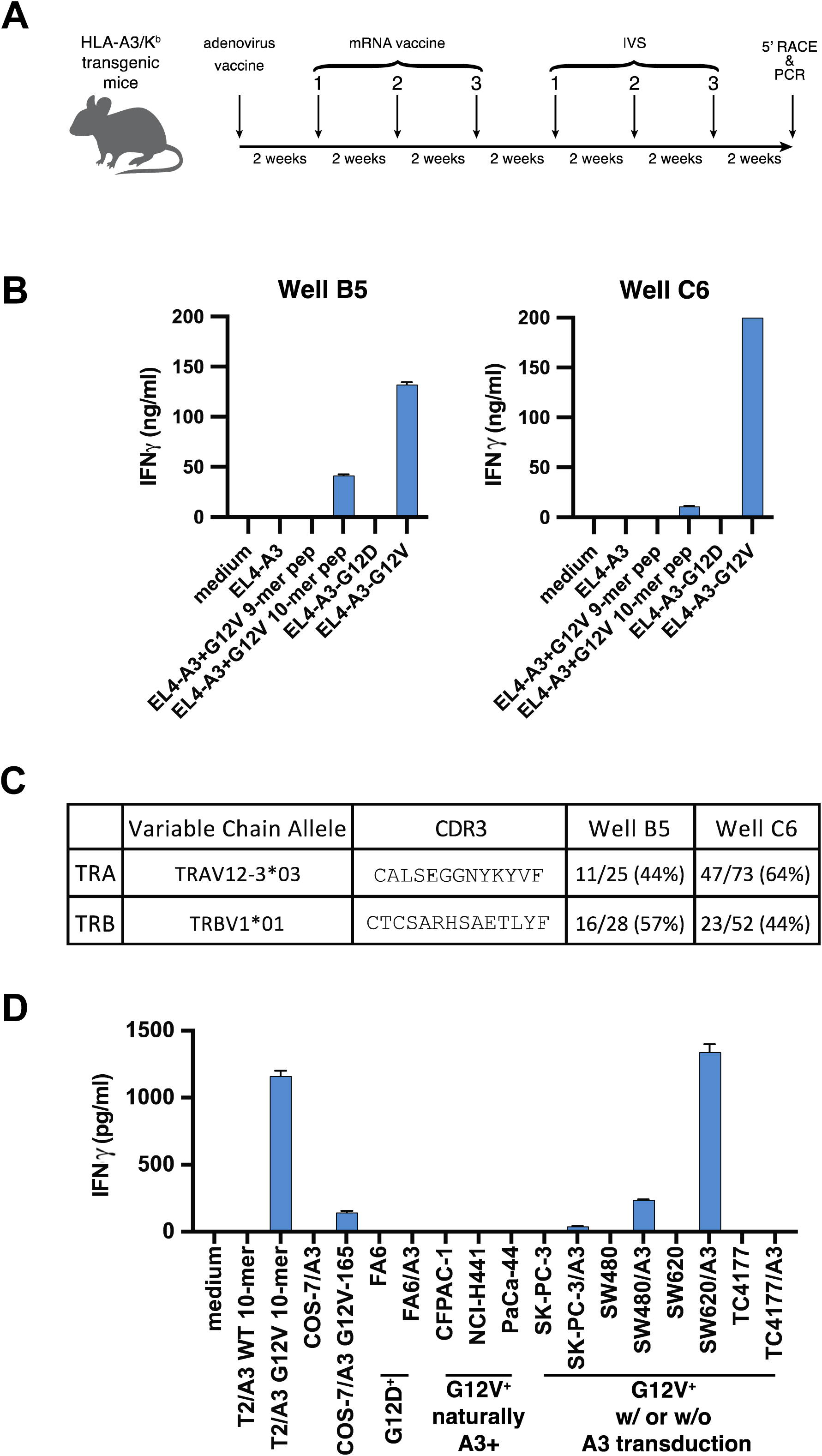
Identification of functional HLA-A3 restricted KRAS-G12V specific TCR by vaccination. (**A**) Vaccination schedule and TCR identification. HLA-A3/Kb transgenic mice (n=2) were vaccinated once with adenovirus encoding KRAS_1-30_, containing the G12V mutation, followed by three vaccinations with mRNA encoding the G12V 10mer epitope sequence (VVVGAVGVGK). All four vaccinations were spaced 2 weeks apart. After 8 weeks, *in* vitro stimulation of splenocytes was performed with 1 mM G12V-10mer or G12V-9mer (VVGAVGVGK) peptides in the presence of IL-2 (30 IU/ml). After three peptide stimulations IFN-g concentration was measured in culture supernatant by ELISA (shown in B). TCR sequences were isolated by 5’ RACE and PCR (shown in C). **(B)** IFN-g release by splenocytes post *in vitro* stimulation: Two wells in a 24-well plate (B5 and C6) from the *in vitro* stimulation showed specificity to KRAS G12V 10-mer and EL4 cells expressing both HLA-A3 and KRAS G12V mutation. **(C)** TCR sequences of cells in reactive wells (B5 and C6) revealed that the most dominant TRA and TRB were shared between these two wells. **(D)** IFN-g release by G12V TCR+ T cells in response to G12V+ HLA-A3+ targets. TCR sequences identified in C were cloned into a retroviral vector and introduced into human PBL. G12V-TCR+ T cells were mixed with various targets to examine HLA-A3 restriction, G12V specificity and sensitivity. For targets loaded with WT or G12V peptides, T2-A3 and COS-7-A3 cells were used. For targets naturally carrying both HLA-A3 and the G12V-KRAS mutation, CFPAC-1, NCI-H441 and PaCa-44 were used. For targets naturally carrying the G12V-mutation but not HLA-A3, SW480, SW620 and TC4177 were used post transfection with HLA-A3. IFN-g concentration was measured in culture supernatant by ELISA.

**Table 1.**
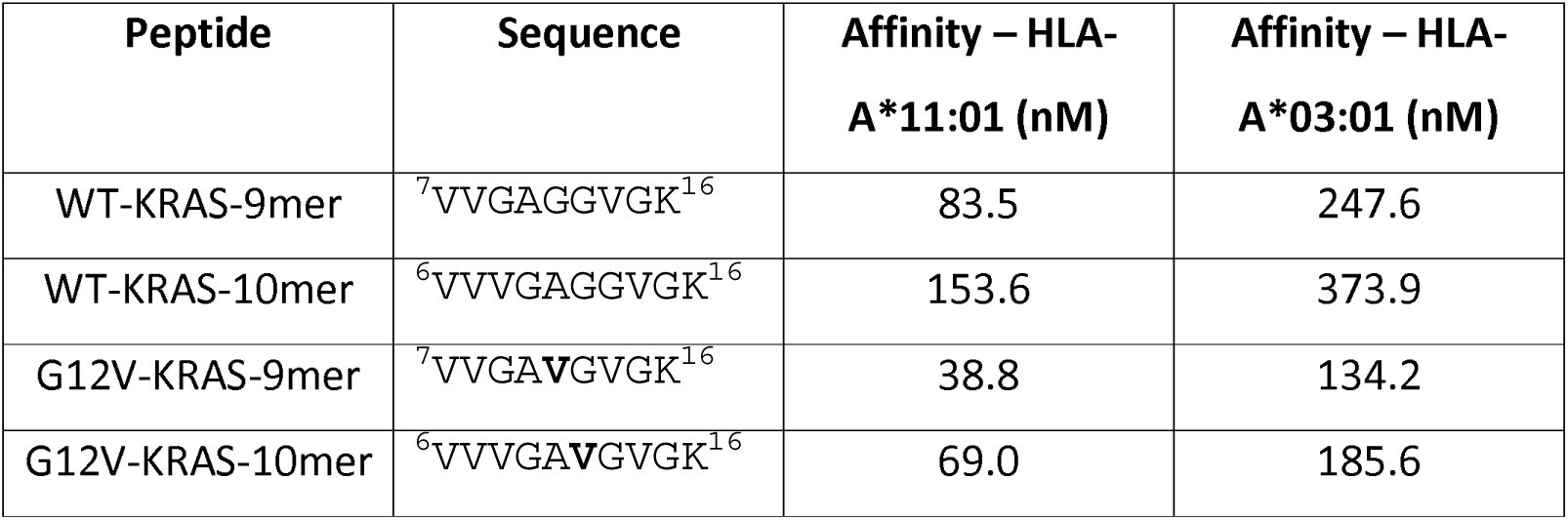
Predicted binding affinities of KRAS peptides to HLA-A11 and HLA-A3.

This TCR was retrovirally transduced into human peripheral blood lymphocytes (PBL) and displayed strong reactivity to HLA-A3^+^ T2 cells loaded with G12V-10mer but not KRAS WT-10mer. Expression of HLA-A3 in tumour cell lines SW480 and SW620 cells, which carry the KRAS-G12V mutation, induced IFN-γ responses from G12V-TCR transduced PBL (Fig. 1D). In contrast, expression of HLA-A3 in FA6 cells, which carry G12D-KRAS, did not induce functional responses. However, G12V-TCR transduced PBL did not respond to tumour lines CFPAC-1, NCI-H441 and Paca-44 that naturally carry both G12V-KRAS and HLA-A3 (Fig. 1D). Thus, we successfully identified G12V-KRAS-10mer specific TCR restricted by HLA-A3, but this TCR may not be sensitive enough to detect G12V-KRAS antigens on certain tumours. Efforts to improve TCR affinity through randomized libraries have led to substantial increases in affinity but also to unwanted and sometimes dangerous off-target effects [21, 22]. We therefore decided to take a structure guided approach to see if we could improve the sensitivity of this TCR through rational substitutions.

### G12V mutation induces an altered peptide conformation bound to HLA-A3

The recombinant soluble HLA-A3 and G12V-TCR were produced as previously described [7, 23, 24] (SFig. 1). We first solved crystal structures of WT and G12V-KRAS 10-mer peptides bound HLA-A3 to resolution of 1.9Å and 2.5Å, respectively (Fig. 2A, Supp. Table 1). Both peptide densities were well defined within the HLA-A3 peptide binding groove (SFig. 2). As expected, the overall structures were very similar with an RMSD of 0.55Å with majority of the peptide-HLA interactions identical between the WT and G12V 10-mer peptide. The major structural difference, beyond the presence of p6Val in G12V-KRAS-A3 complex, was the orientation of p5 Ala which flipped 95° from facing the α1 helix in the WT peptide to facing the α2 helix in the G12V peptide complexed structures (Fig. 2B-E). This results in p5Ala making two van der waals (VDW) contacts with Q66 and A69 of HLA by the WT peptide but only one contact with Q155 by the G12V peptide (Fig. 2D, E). However, despite the fewer contacts made by G12V-KRAS-10mer with HLA-A3 (Extended Data Table 1), the mutant peptide displayed a slightly higher melting temperature than WT-KRAS (Fig. 2F).

**Figure. 2.**
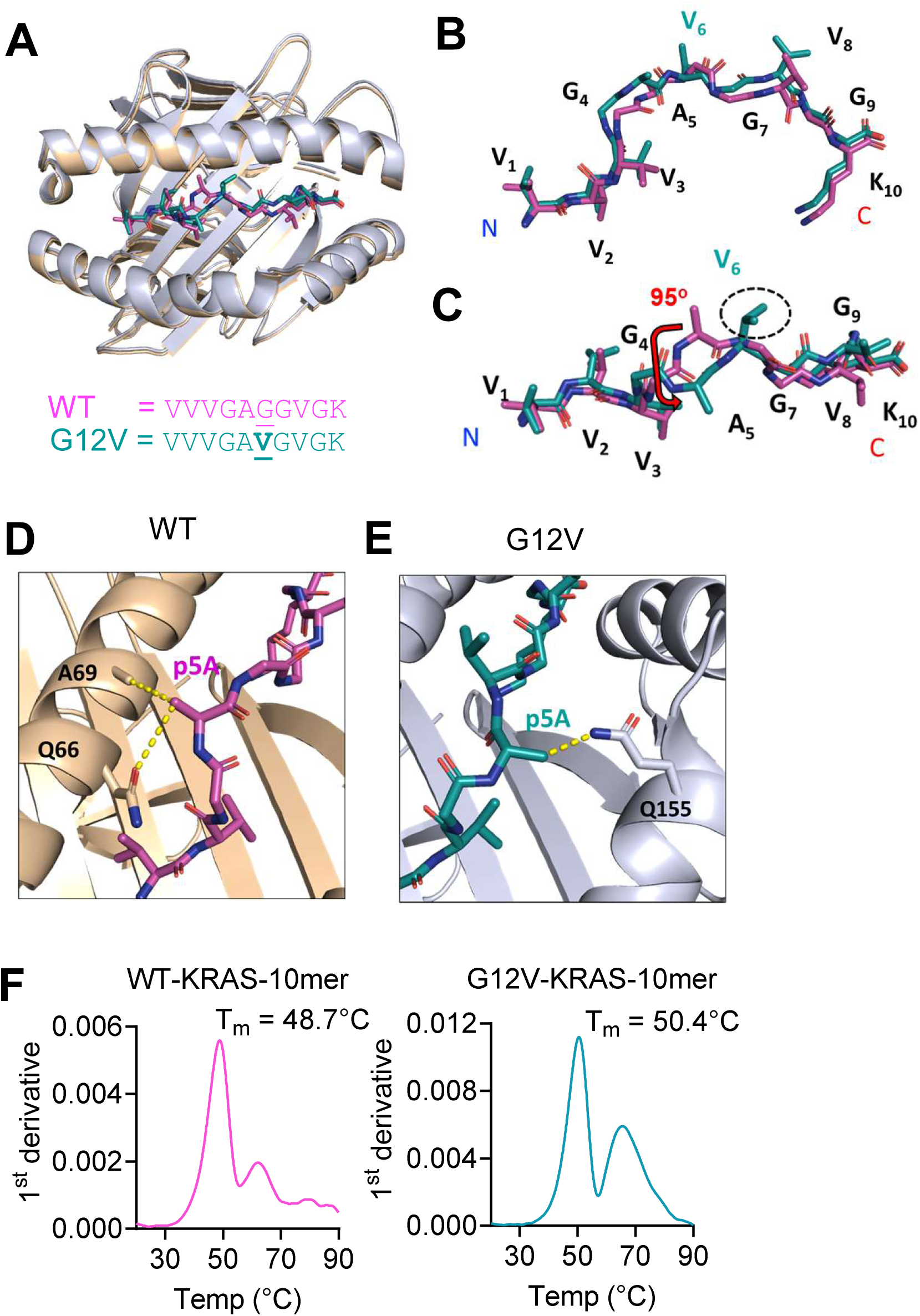
Altered conformation of mutant G12V peptide presented by HLA-A*03:01. (**A**) Overlay of HLA-A*03:01 structures bound to KRAS-WT (pink) and mutant KRAS-G12V (turquoise) peptides. **(B,C)**. Side **(B)** and top **(C)** view of WT and mutant G12V-KRAS peptides. Position of p6 Val (G12V-mutant) and p5 Ala rotation is noted. **(D)** Interactions of HLA-A*03:01 a_1_ helix with p5 Ala of WT KRAS peptide**. (E)** Interactions HLA-A*03:01 a_2_ helix in mutant G12V-KRAS with p5 Ala of mutant G12V-KRAS peptide. **(F)** Thermal melting curves for HLA-A3 bound to WT and G12V KRAS-10mers, n = 2. PDB codes: 8VJZ (WT) and 8RNI (G12V).

### Specificity and affinity of G12V-TCR

We next determined the affinity of this TCR by surface plasmon resonance (SPR). We observed no detectable binding of G12V-TCR to HLA-A3 refolded with WT-10mer (WT-A3) with TCR concentrations up to 40 μM (Fig. 3A). In contrast, G12V-TCR bound to HLA-A3 refolded with G12V-10mer (G12V-A3) with an affinity (K_D_) of 23 μM (Fig. 3B, C). While G12V-TCR exhibits specificity to the KRAS neoantigen, the binding affinity is weaker than that observed with therapeutic KRAS-G12D specific TCRs defined previously [2, 5, 7].

**Figure. 3.**
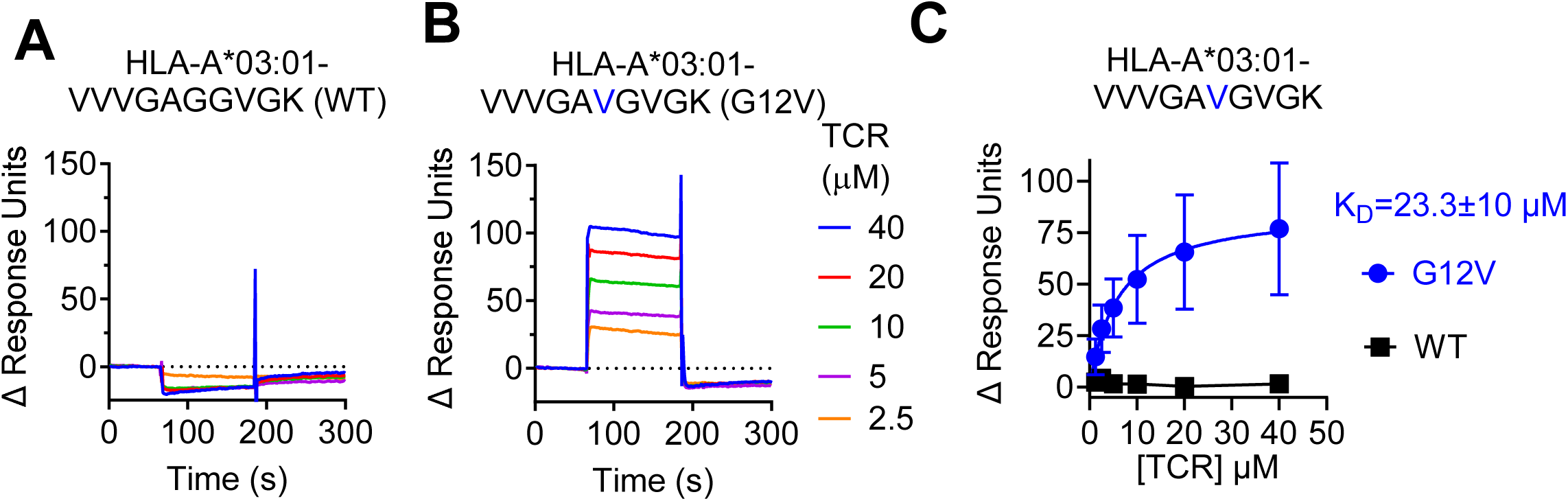
Specificity and affinity of mutant KRAS-G12V TCR. (**A, B**) Representative SPR sensorgrams displaying G12V-TCR binding to captured HLA-A*03:01 refolded with b2-microglobulin and the WT KRAS peptide (VVVGAGGVGK) **(A)** and the mutant-G12V KRAS peptide (VVVGA**V**GVGK) **(B)**. Binding to reference flow cell is subtracted. **(C)** Equilibrium analysis of SPR measurements of G12V-TCR binding shown in **(A)** and **(B)**. Mean responses at each analyte concentration and standard error between n=3 injections (error bars) are shown. Non-linear regression curve fit of the one-to-one specific binding model (line) are shown. K_D_ value ± standard deviation is shown calculated by modelling steady state kinetics. No K_D_ was determined for G12V-TCR binding to WT peptide.

### Structure of G12V TCR in complex with G12V-HLA-A3

To understand how G12V-TCR discriminates between the WT and G12V peptide bound HLA-A3, we solved structures of the receptor alone as well as its complex with G12V-A3 to 1.7Å and 3.5Å resolutions, respectively (Fig. 4A, SFig. 3, Supp. Table 1). There were eight TCR-G12V-A3 complexes in the crystallographic asymmetric unit with highly similar structures (SFig. 3A&B average RMSD to complex efghi = 0.434 ± 0.102). The structure of the CDR loops were very similar between the TCR alone and TCR bound structures (SFig. 3C&D). TCR recognition was dominated by the receptor beta chain with more than twice as many contacts as the alpha chain (57 versus 26) (Fig. 4B, Extended Data Table 1). Highlighting this β chain dominance, none of the TCRα CDRs contacted the peptide and CDR2α made no contact to the HLA either. The four TCRα residues that contacted the HLA-A3 α1 helix are Y30 of CDR1α and G97, N99 and Y100 of CDR3α (Fig. 4C). In contrast, TCRβ CDR1, 2 and 3 made contacts to HLA-A3 with R54 and S55 of CDR2β interacting with the HLA-A3 α1 helix (Fig. 4D), and Q32 of CDR1β, and R99, E103 of CDR3β interacting with the HLA-A3 α2 helix (Fig. 4E).

**Figure. 4.**
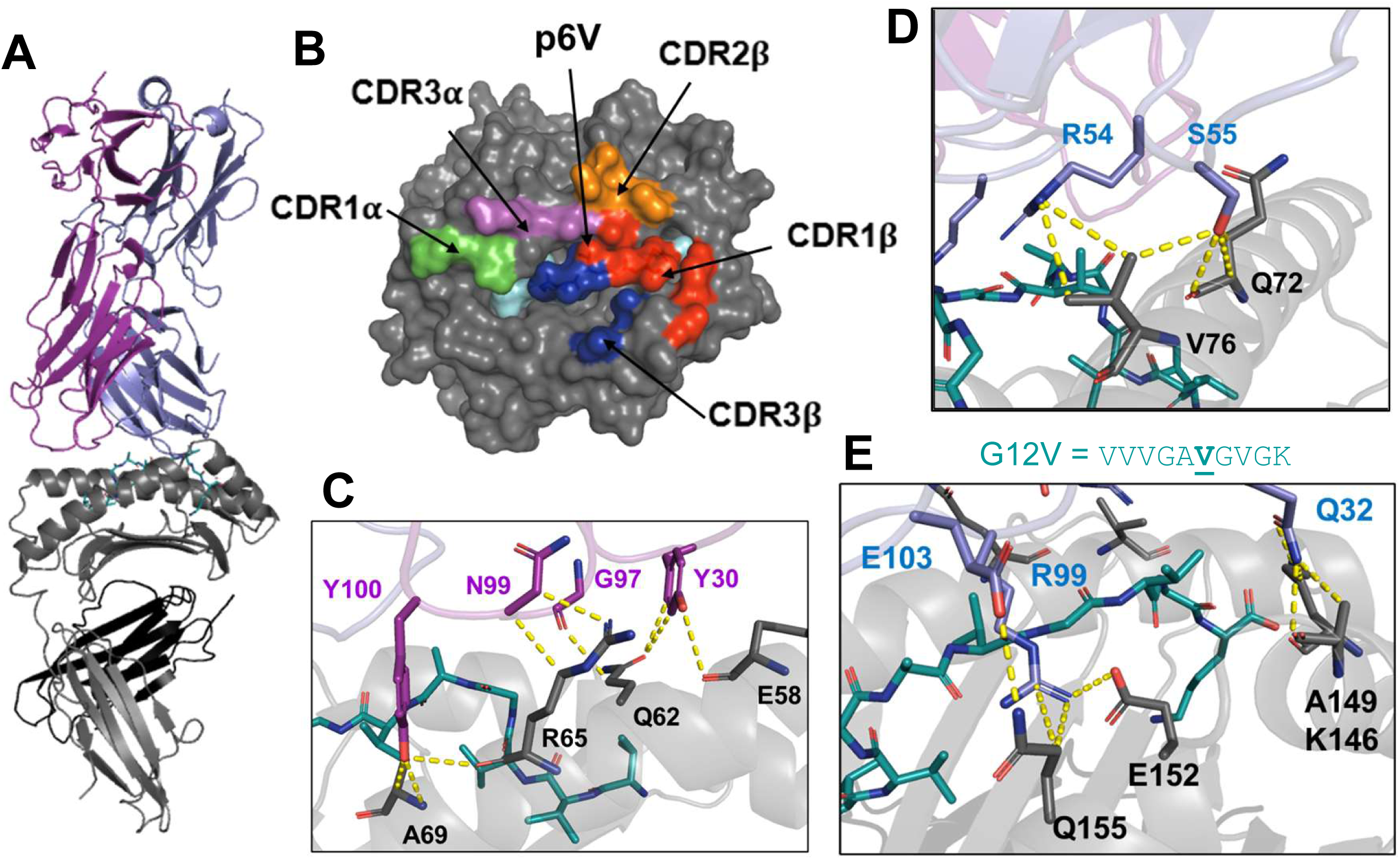
Structure of G12V-TCR in complex with mutant G12V-KRAS bound to HLA-A*03:01. (**A**) Overall structure of G12V TCR (a chain; purple, b chain; blue) with HLA-A*03:01 (grey) and bound mutant G12V-KRAS peptide (turquoise). **(B)**. Location of G12V-TCR footprints on structure of mutant G12V-KRAS peptide bound to HLA-A*03:01. Alpha chain CDR1 (green), CDR3 (pink), Beta chain CDR1 (red), CDR2 (orange) and CDR3 (blue). **(C)** Contacts between G12V-TCR alpha chain (purple) and HLA-A*03:01 amino acids (dark grey). **(D, E)** Contacts between G12V-TCR beta chain (blue) and HLA-A*03:01 amino acids (dark grey). G12V-10mer peptide is shown in turquoise.

### G12V-TCR recognizes G12V mutation via a TCRβ hydrophobic pocket

Val residue at p6 of the G12V peptide interacts W35 from CDR1β and H100 from CDR3β of G12V-TCR with both CDR residues making multiple VDW contacts to the sidechain of p6Val (Fig. 5A, SFig. 3E). These interactions are predicted to be absent in the WT-KRAS peptide with p6Gly bounded HLA-A3 (Fig. 5B-C), suggesting they form the structural basis for antigen recognition and discrimination between WT-A3 and G12V-A3. We next developed a tetramer binding assay to assess the contribution of different TCR residues to specific recognition of G12V-A3 (SFig. 5). We generated 293T cells expressing murine CD3 that were subsequently transfected with unmodified G12V-TCR or variants with substitutions in key TCR contacts. Consistent with functional and SPR data (Fig. 1&3), HLA-A3 tetramers with G12V but not WT peptide bound to G12V-TCR (Fig. 5D). Individual substitutions, either W35A on CDR1β or H100A on CDR3β completely abolished binding (Fig. 5D), further demonstrating their importance in G12V mutant antigen recognition. The CDR3α Y100A substitution also reduced G12V tetramer binding but was less severe than the CDR1/3β substitutions, highlighting the beta chain dominance of this TCR. These data provide a strong structural rationale for how G12V-TCR discriminates between WT and G12V-KRAS 10mer peptides, underpinning its potential to target tumours. Consistently, a recent study defined the structural basis for detection of G12V-A3 by a peptide specific mAb to also be dominated by hydrophobic interactions [19]. Here, the heavy chain CDR3 and light chain CDR2 form a hydrophobic pocket surrounding p6Val, with Ile102, Pro103 and Val103 of the CDR3^H^ forming VDW contacts, in addition to peptide conformational changes not observed in our structures [19]. This convergent structural mechanism for detecting G12V-A3 suggests that targeting this neoantigen via other TCRs of mAbs will require a similar hydrophobic dependent mechanism.

**Figure. 5.**
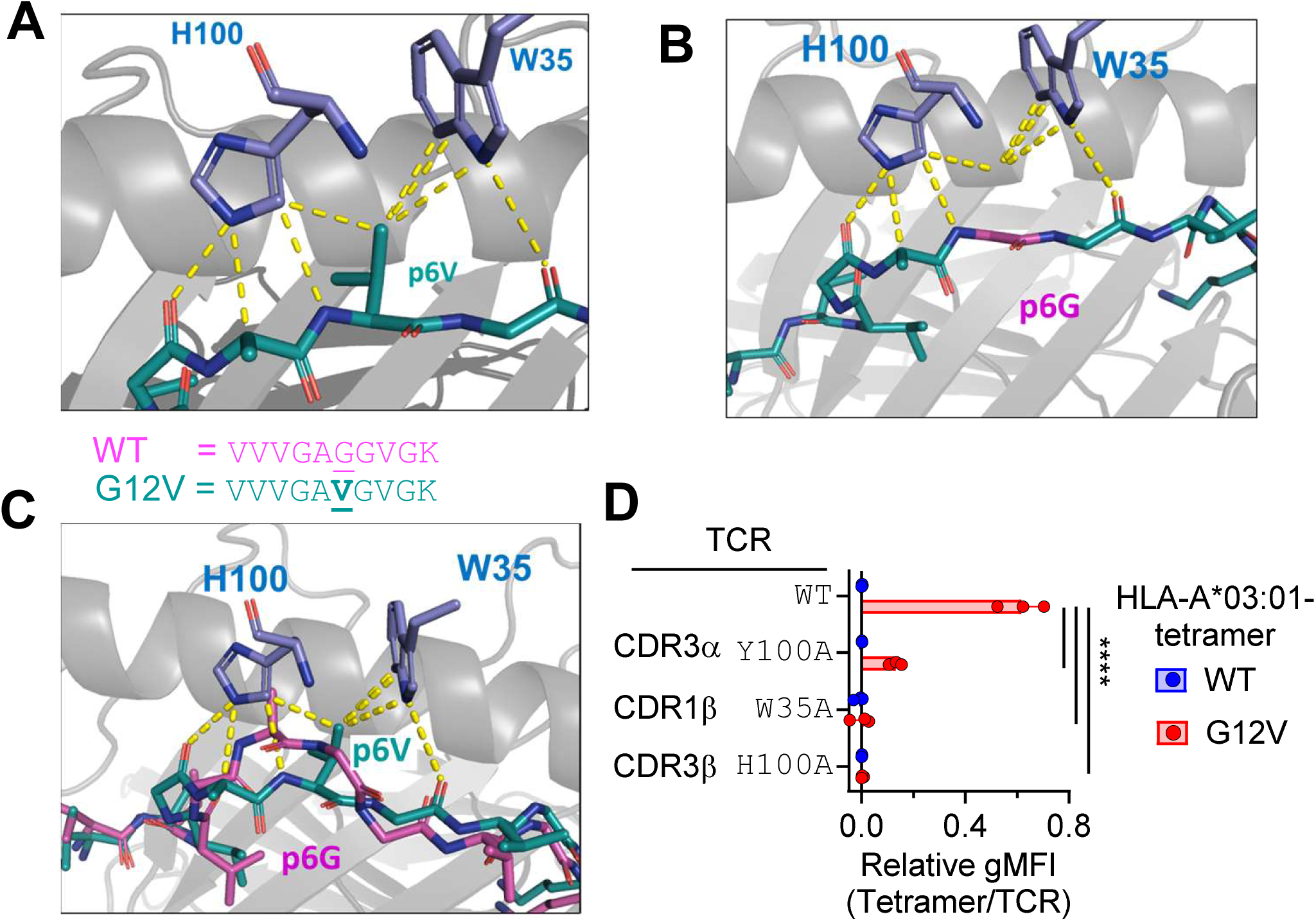
Molecular basis for TCR specificity to mutant G12V peptide. (**A**) Interactions between CDR1b W35, CDR3b H100 (blue) and mutant G12V-KRAS peptide (turquoise). **(B)** Modelling G12V-TCR CDR1b W35, CDR3b H100 interactions with WT KRAS peptide by modelling p6 Val – Gly. **(C)** Modelling G12V-TCR CDR1b W35, CDR3b H100 interactions with WT and mutant G12V-KRAS peptides by overlaying structure of WT KRAS peptide (pink) shown in Fig. 4. **(D)** Impact of indicated substitutions in G12V-TCR on binding to HLA-A*03:01 tetramers refolded with WT (blue) and mutant G12V-KRAS peptides (red). Binding was measured by flow cytometry (n = 3) and normalised to TCR expression. **** = p<0.0001 by student t-test (two tailed).

### Rational substitutions modestly improve G12V-TCR binding to G12V-KRAS-A3

While G12V-TCR displayed specificity for KRAS-G12V-10mer, T cells expressing this receptor were not stimulated by tumours naturally expressing HLA-A3 and carrying the KRAS-G12V mutation (Fig. 1). There were no obvious rational substitutions to improve interactions between G12V-TCR and peptide p6Val and therefore we decided to make TCR substitutions improving interactions with the HLA-A3 heavy chain. We modelled 7 substitutions in the TCR CDR1α and CDR2β to modify interactions with HLA-A3 (Fig. 6A, SFig. 5) and tested these substitutions for improving TCR-HLA-A3 interactions using the tetramer binding assay described earlier. Only one substitution (CDR1α I29Q) enhanced TCR binding to WT-A3 tetramers, suggesting this approach would not compromise the tumour specificity of this TCR. Four substitutions on CDR1α, I29H, I29Q, S31D, and S31E had no effect or slightly decreased TCR binding to G12V-A3 tetramers, while one CDR2β substitution, S55D, abolished tetramer binding (Fig. 6A). Only CDR1α I29K and CDR2β D58E showed mild increases in the tetramer binding and the improvement was consistent across multiple tetramer concentrations (Fig. 6A-B).

**Figure. 6.**
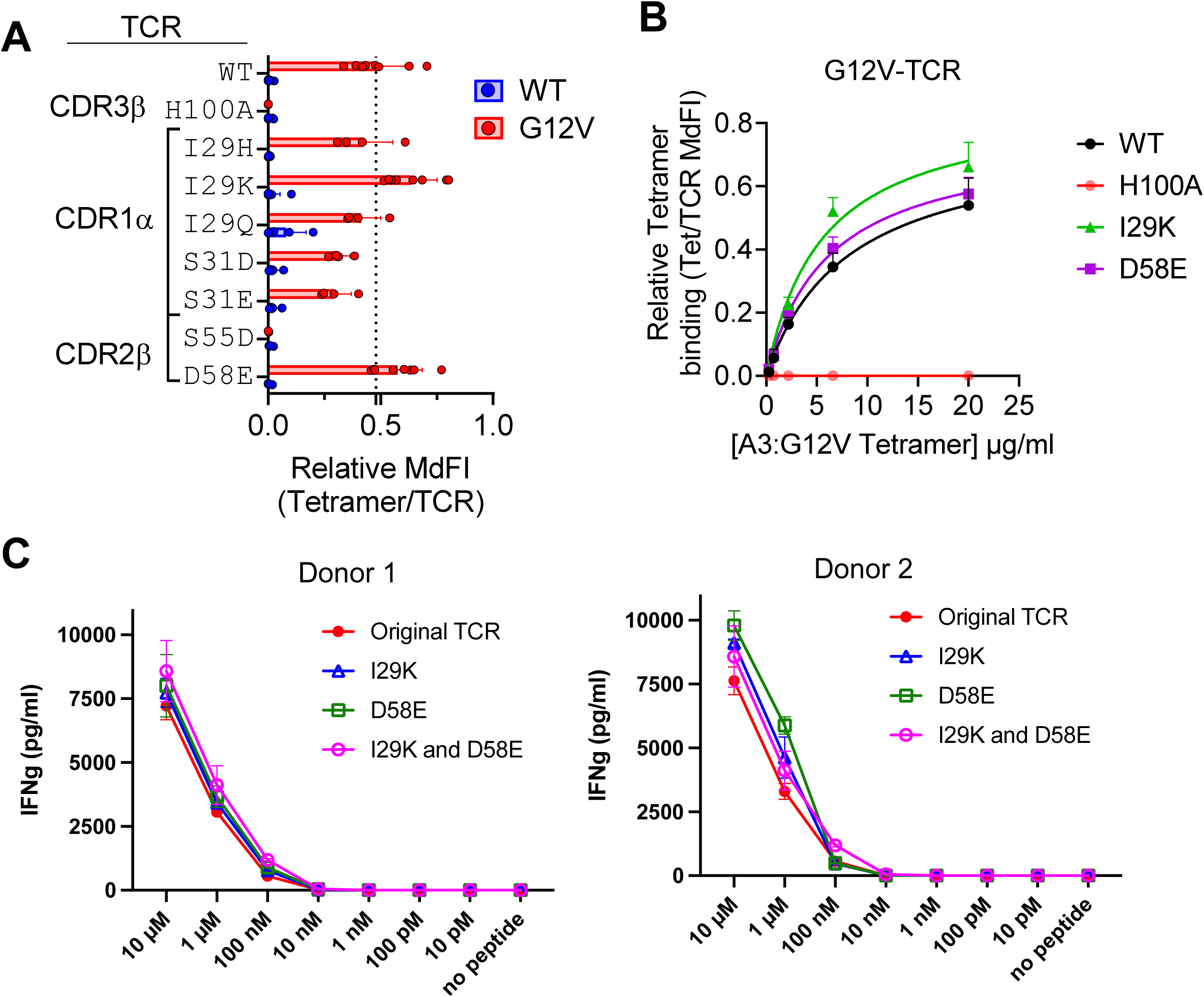
Rational substitutions modestly improve G12V-TCR binding. (**A**) Impact of indicated substitutions in G12V-TCR on binding to HLA-A*03:01 tetramers refolded with WT (blue) and mutant G12V-KRAS peptides (red). Binding was measured by flow cytometry (n = 4-8) and normalised to TCR expression. **(B)** Titration of HLA-A*03:01-G12V-KRAS binding to G12V-TCR with indicated substitutions (n=3). **(C)** IFN-g production by human T cells expressing G12V-TCRs with indicated substitutions in response to COS-7 cells expressing HLA-A*03:01 preloaded with mutant G12V-KRAS peptide. Data from two donors (left and right) are shown. IFN-g was measured by ELISA.

To investigate if the improved G12V-A3 binding of the I29K and D58E mutant TCR resulted in enhanced T cell function, we measured IFN-γ secretion from original, I29K, D58E or double substituted G12V-TCR transduced PBL after stimulation with the G12V-10mer pulsed HLA-A3+ COS-7 cells (Fig. 6C, SFig. 6). Both the single and double substituted TCR transduced PBL exhibited a modest but consistent increase in total IFN-γ secretion at 10 μM – 100 nM concentrations of peptide than the WT G12V-TCR transduced cells. However, none of the substitutions improved TCR sensitivity to lower concentrations of peptide (<1nM) suggesting they may not be effective in clinical settings.

## Discussion

Public neoantigens are attractive targets for precision medicines as they can target the same mutation in many patients [2, 3]. However, patient eligibility is limited by HLA polymorphism, which determines whether a given neoantigen peptide can be displayed by a tumour [2–4]. Therefore, it is imperative to identify candidate antigens and TCRs across as many common HLA types as possible. Here we identified a murine TCR specific for the KRAS-G12V mutation restricted by the common HLA-I allotype HLA-A3 using a vaccination approach in HLA-A3 transgenic mice. We determined crystal structures of HLA-A3 presenting WT-KRAS and G12V-KRAS peptides demonstrating a conformational change in the G12V-KRAS peptide. We determined the solution binding affinity of this TCR, and its structural mechanism to discriminate WT– and G12V-KRAS peptides. Using a structure guided approach, we designed and tested substitutions predicted to increase TCR-HLA binding and identified two substitutions that marginally increased TCR binding and function. However, these substitutions exhibited no improvement at clinically relevant concentrations of the peptide antigen.

Neoantigens fall into two broad categories, type 1 and type 2 [2, 8]. Type 1 neoantigens contain substitutions in existing HLA bound peptides, while type 2 neoantigens are generated by the substitution forming a previously absent critical anchor residue [2, 8]. Our previous work on HLA-C*08:02 restricted KRAS-G12D specific TCRs, demonstrated these to be type 2 neoantigens, where the G12D mutation formed a critical anchor residue necessary for HLA-C*08:02 stabilization [7]. The G12V-KRAS 10mer presented by HLA-A3 is a type 1 neoantigen, with the substitution situated within an existing ‘self’ epitope. Intuitively, the G12V-TCR focuses on the substitution site, utilizing a hydrophobic pocket generated by CDR1β W35 and CDR3β H100 to accommodate the p6Val of G12V-KRAS-10mer. This dependence of hydrophobic interactions for detection of G12V-A3 is consistent with a recent study of a peptide-specific mAb that also forms hydrophobic interactions with p6 Val [19]. This mode of detecting type 1 neoantigens has been observed previously, such as by HLA-A*02:01 restricted TCRs specific for p53 R175H, although other modes for detecting type 1 neoantigens have been described [2, 25, 26].

Considerable efforts have been made to identify TCRs specific for public neoantigens for the development of multiple therapeutic options including TCR-T and peptide-specific biologicals [3, 5, 9, 11–13, 15, 19, 20]. The G12V-KRAS specific TCR identified here was highly specific for the mutant KRAS peptide and not WT, but tumours carrying both HLA-A3 and KRAS-G12V did not stimulate this TCR when expressed in primary human T cells. Recent studies have identified both TCRs and single-chain variable fragments (scFv) that can detect G12V-KRAS-10mer when presented on HLA-A3+ tumour cells with anti-tumour efficacy *in vivo* [11, 20]. It is estimated that only 9 copies of G12V-KRAS-10mer are found per cell on the HLA-A3+ tumour line NCI-H441 [20], suggesting our TCR failed due to an inability to detect so few ligands, perhaps due to its relatively low affinity of 22 μM [2]. However, the affinity of the TCR identified by Bear et al. [11] is unknown, preventing a direct comparison. Our recent work on KRAS-G12D specific TCRs suggests that TCRs with affinities in the high nM to low μM range are most effective for anti-tumour function, in line with a ‘goldilocks’ zone for TCR affinities [2, 7, 27, 28]. However, other models have suggested that low avidity TCRs are superior to higher avidity [29]. More investigations into the affinities of effective anti-tumour TCRs are required to understand the desired biophysical properties of TCRs to be used in the clinic.

Finally, while this HLA-A3 restricted G12V-KRAS TCR is not sensitive enough to detect naturally occurring tumors, it provides a platform for further development of high affinity TCRs through library-based affinity selection [13]. While it is necessary to identify additional candidate TCRs for clinical development, understanding the functional, biophysical and structural properties of TCRs that do not progress to the clinic can provide important benchmarks to understand efficacious anti-tumour responses and effective T cell responses more generally [30].

## Materials and Methods

### Animal Experiments

This study followed the NIH Animal Care and User Committee (ACUC) guidelines, animal protocol number SB201.

### Cell culture medium

D10 (DMEM with 10% FBS), R10 (RPMI1640 with 10% FBS), CM (RPMI1640 supplemented with 10% FBS, Non-Essential Amino Acid Solution, 1mM Sodium Pyruvate, 55 µM 2-Mercaptoehanol), RH10 (RPMI1640 with 10% heat-inactivated human AB serum (Valley Biomedical). RH10-300 (RH10 medium supplemented with 300 IU/ml recombinant human Interleukin-2).

### Cell lines

Cell lines cultured in D10: HEK293T cells (RRID: CVCL_0063), 293GP (RRID: CVCL_E072), COS-7 (RRID: CVCL_0224), FA6 (RRID: CVCL_4034), CFPAC-1 (RRID: CVVL_1119), NCI-H441 (RRID: CVCL_1561), PaCa-44 (RRID: CVCL_7087), SK-PC-3 (RRID: CVCL_4055), SW480 (RRID: CVCL_0546), SW620 (RRID:CVCL_0547). Cell lines cultured in R10: T2 (RRID: CVCL_2211) and its derivatives. Cell lines cultured in CM: EL4 (RRID: CVCL_0255) and its derivatives.

### Peripheral blood lymphocytes from normal donors

Peripheral blood lymphocytes (PBL) were obtained from healthy donors who provided informed consent for the utilization of their lymphocytes in research, following the clinical protocol NCT00068003.

### Murine vaccination and identification of G12V-KRAS specific TCR

An adenovirus plasmid encoding the first 30 amino acids of KRAS with the G12V mutation (KRAS G12V-30) was created following the manufacturer’s instructions (Adeno-X CMV, Takara Bio). Adenovirus particles were produced by transfecting the adenoviral plasmid into HEK293 cells. The virus particles were then purified using the Aneno-X Maxi Purification Kit (Takara Bio), and the buffer was replaced with 1X Formulation Buffer containing 2.5% glycerol (w/v), 25 mM NaCl, and 20 mM Tris-HCl pH8.0. The virus titer was measured using Adeno-X Rapid Titer Kit (Takara Bio) and 10^8^ pfu (plaque-forming units) of adenovirus were injected intravenously into C57BL6/HLA-A3-K^b^ transgenic mice. The mice were subsequently immunized three more times with intra-muscular injections of 10 µg of mRNA encoding the sequence MVVVGAVGVGK, and 12.5 µg of helper RNA encoding MNNRKWFPAEPEDVRDYLLYLQARFFLETEETFQPGRWFMRAAQAVTAVVCGPDMIQVSLRDQRPVGIRPRSRYLTAAAVTAVLQDFADIFLMQNMAEFDDQLIFNSISARALKAYFTAKINEMVDGVITEQQSSINISGYNFSLGAAVKAGAALLDGGNM (Moderna) for each immunization. Two weeks after the final immunization, splenocytes were harvested and subjected to in vitro stimulation. Stimulation was performed using a 9-mer peptide (VVGAVGVGK) or a 10-mer (VVVGAVGVGK) peptide at a concentration of 1 µM. This stimulation took place in 24-well plates with CM with recombinant human interleukin 2 (IL-2) 30 IU/ml. After two weeks of stimulation, T cells were further stimulated with lymphoblasts treated with ConA and pulsed with the cognate peptide (1 µM). Subsequently, 2 weeks later, T cells underwent further stimulation using EL4 cells transduced with retrovirus encoding HLA-A3-K^b^ and KRAS G12V-30. Following this stimulation, the culture was assessed for reactivity, including reactivity to EL4/A3-Kb, EL4/A3-Kb pulsed with the cognate peptide, and EL4/A3-Kb-KRAS G12V-30. Total RNA was extracted from cells in two wells exhibiting reactivity to EL4/A3Kb-KRAS G12V-30. The TCR sequences were then isolated using 5’RACE (SMARTer RACE cDNA Amplification Kit, Takara Bio), followed by PCR amplification with primers designed to be complementary to TRAC (GTTGCTCCAGGCAATGGCCCCATTGCTC) and TRBC (GGTCCGTGCTGACCCCACTGTGGACCTC) following the manufacturer’s protocol.

### TCR transduction of PBL

The T cell receptor alpha (TRA) and beta (TRB) genes underwent codon optimization to enhance their expression in human cells. Subsequently, these optimized genes were cloned into the retroviral vector pMSGV1 in the following order: TRB, P2A, and TRA. Retroviral transduction of TCR genes were recently described [31]. Briefly, on day 1, 5×10^6^ 293GP cells were seeded onto poly-D-Lysine-coated 10 cm dishes in D10. On day 2, these cells were transfected with the retroviral plasmid and the RD114 envelope plasmid (retroviral plasmid 25 µg, RD114 plasmid 3 µg, and 70 µl of Lipofectamine 2000). Simultaneously on day 2, human peripheral blood cells (2×10^6^ cells/well in a 24-well plate) were stimulated with anti-CD3ε antibody (OKT3) at a concentration of 200 ng/ml in RH10-300 medium. On day 3, D10 medium for 293GP was refreshed. On day 4, the 293GP culture supernatants were filtered through a 0.45 µm syringe filter and loaded onto Retronectin (Takara Bio)-coated 24-well plates (1 ml per well) by centrifugation at 2,000G, 32°C for 2 hours. Subsequently, the OKT3-stimulated T cells suspended in RH10-300 medium were added to the virus-loaded wells at a density of 5×10^5^ cells/ml/well and centrifuged again at 1,000G for 10 minutes.

### T cell functional assays

TCR transduction efficiency analysis by flow cytometry was performed using anti-human CD3 APC-H7 (BD Biosciences #641406), anti-human CD8 PE-Cy7 (BD Biosciences #335787), and anti-mouse TCR β chain FITC (BD Biosciences #H57-597). For the IFN-γ secretion assay, COS-7 cells retrovirally transduced with HLA-A*03:01 gene were pulsed with the denoted concentration of peptides in R10 for 2 hours at room temperature. The cells were then washed three times, and 5×10^4^ cells/well in a 96-well flat-bottom plate were cocultured with 1×10^5^ TCR-transduced cells/well, suspended in R10. After 20 hours, IFN-γ in the culture supernatant was measured by ELISA (DuoSet, R&D Systems).

### Recombinant protein production

Recombinant G12V-TCR and HLA-A3 were produced by *in vitro* refolding of dissolved inclusion bodies expressed in bacteria, largely as described [7, 24, 32]. DNA encoding HLA-A3 (residues 1-274), β_2_M (1-99), G12V-TCR alpha chain (1-206) and G12V-TCR beta chain (1-246) were synthesized by Genscript (USA) and cloned into the pET30a+ vector. Proteins were expressed in *E. coli* (strain BL21-DE3, Invitrogen, USA) by growing from antibiotic resistant colonies. Bacteria were lysed and inclusion bodies washed before dissolving in 8 M urea, 20 mM Tris (pH 8.0), 0.5 mM EDTA and 1 mM DTT. Dissolved inclusion bodies were added dropwise to cold refolding buffer before dialysis and purification. Refolding buffer was 1 L of 0.1 M Tris (pH 8.0), 0.4 M Arginine, 2.5 mM EDTA, 0.5 mM oxidized glutathione and 5 mM reduced glutathione. 50mg of HLA-A3 heavy chain, 10 mg of β2m, A3-peptide-β_2_M complexes were refolded by adding 10 mg of peptide (Genscript USA, dissolved in DMSO). TCR was refolded by adding 30 mg of TCRα and TCRβ into 1L refolding buffer containing 0.4 Arginine, 0.1M Tris (pH 8.0), 2.5 mM EDTA, 0.5 mM oxidized glutathione and 5 mM reduced glutathione. Proteins were dialyzed against 10 mM Tris (pH 8) and purified by ion exchange (Q column) followed by size exclusion (S200, GE Healthcare) chromatography. Protein fractions were assessed for purity by SDS PAGE, pooled and concentrated.

### Surface plasmon resonance

SPR was performed largely as described [7, 33]. SPR binding experiments were performed with a BIAcore 3000 instrument and analyzed with BIAevaluation software v4.1 (GE Healthcare). CM5 chips (GE Healthcare) were coated in the HLA-I–specific mAb W6/32 (BioLegend) by primary amine coupling at 5,000 to 7,000 response units (RUs) using a 2 μL/min flow rate in 10 mM sodium acetate (pH 5.5). HLA-A3 was captured by W6/32 at approximately 600 RUs in PBS. The analytes were G12V-TCR heterodimers in 10 mM Hepes (pH 7.5) and 0.15 M NaCl with a flow rate of 50 μL/min. TCRs were injected for 2 min followed by a dissociation of 10 min. Binding was measured with serial dilutions of G12V-TCR from 40 to 1.25 μM. Dissociation constants were obtained by modeling steady-state kinetics with BIAevaluation software.

### Differential scanning fluorimetry (DSF) thermal stability assay

The thermal stability of the HLA-A*03:01 peptide complexes was measured by differential scanning fluorimetry (DSF). The HLA-A*03:01 peptide complexes were prepared at a concentration of 1.0 mg/ml in PBS buffer with a final volume of 20 μL. Each 20 μL sample was split between two Prometheus Grade Standard Capillaries (Nanotemper, Munich, Germany) and placed in the capillary tray of a Prometheus Panta (Nanotemper) instrument. Excitation power was adjusted to cover a range of 8000 to 15,000 Raw Fluorescence Units, with fluorescence emission detection at 330 nm and 350 nm. A 1°C/min thermal ramp over a range of 20°C to 90°C, was applied. Automated thermal melt data calling was achieved using the Thermal Unfolding Analysis software within PR.Panta Analysis, v1.2.

### Crystallography, data collection, structure determination and refinement

Proteins were concentrated to 10 mg/ml before screening for crystallization conditions using a Crystal Gryphon (ARI). WT-KRAS-A3 and G12V-KRAS-A3 crystallized in 0.1M ammonium citrate tribasic pH7, 12% PEG 3350. TCR-G12V-KRAS-A3 complex crystallized in 0.1M Bis-Tris Propane (pH 6.5), 0.2M Na_2_SO_4_, 14-20% PEG 3350. Crystals were coated in cryoprotectant (condition plus 20% Glycerol) before flash freezing in liquid N_2_. All data were collected on the SER-CAT 22 BM or ID beamlines at the Argonne National Laboratory, Illinois, USA. Data were processed and merged with HKL2000 [34] and structures determined by molecular replacement with Phaser in CCP4 [35, 36]. Models were refined with Coot and Phenix [37, 38]. Search models were the HLA-A3 structure (PDB: 2XPG)[39] with the peptide omitted and a mVβ1*01 containing TCR (PDB: 6DFS). Solved HLA-A3 and G12V-TCR structures were used as search models for the TCR-A3(G12V) complex.

### Tetramer binding assay

HEK-293T cells were transfected via Lipofectamine 2000 (Invitrogen, USA) with a plasmid encoding murine CD3δγεζ and FACS sorted on GFP expression. The murine CD3 WTdelta-F2A-gamma-T2A-epsilon-P2A-zeta pMIA II plasmid was a gift from Dario Vignali (Addgene plasmid # 52093) [40]. A plasmid encoding full length G12V-TCR in the format TCRα-T2A-TCRβ was synthesized by Genscript and cloned into pcDNA3.1. Amino acid substitutions in G12V-TCR were generated by quick-change site-directed PCR mutagenesis (Agilent, USA). TCRs were transiently transfected into 293T-mCD3-GFP cells and stained separately with PE anti-mTCRβ mAb (Biolegend, 109208) and HLA-A3 tetramers refolded with WT– and G12V-KRAS 10mer peptides. Biotinylated HLA-A3 monomers were generated and conjugated to streptavidin PE (Agilent, PJRS25-1) as described [41].

### Data deposition

TCR sequences are available in GenBank with accession numbers OR751275 for TRA and OR751276 for TRB. Crystallographic data have been deposited in the Protein Database with the following codes; HLA-A3-WT-KRAS: 8VJZ, HLA-A3-G12V-KRAS: 8RNI, G12V-TCR: 8RO5 and G12V-TCR-HLA-A3(G12V) complex: 8RRO.

## Fig

**SFigure. 1.**
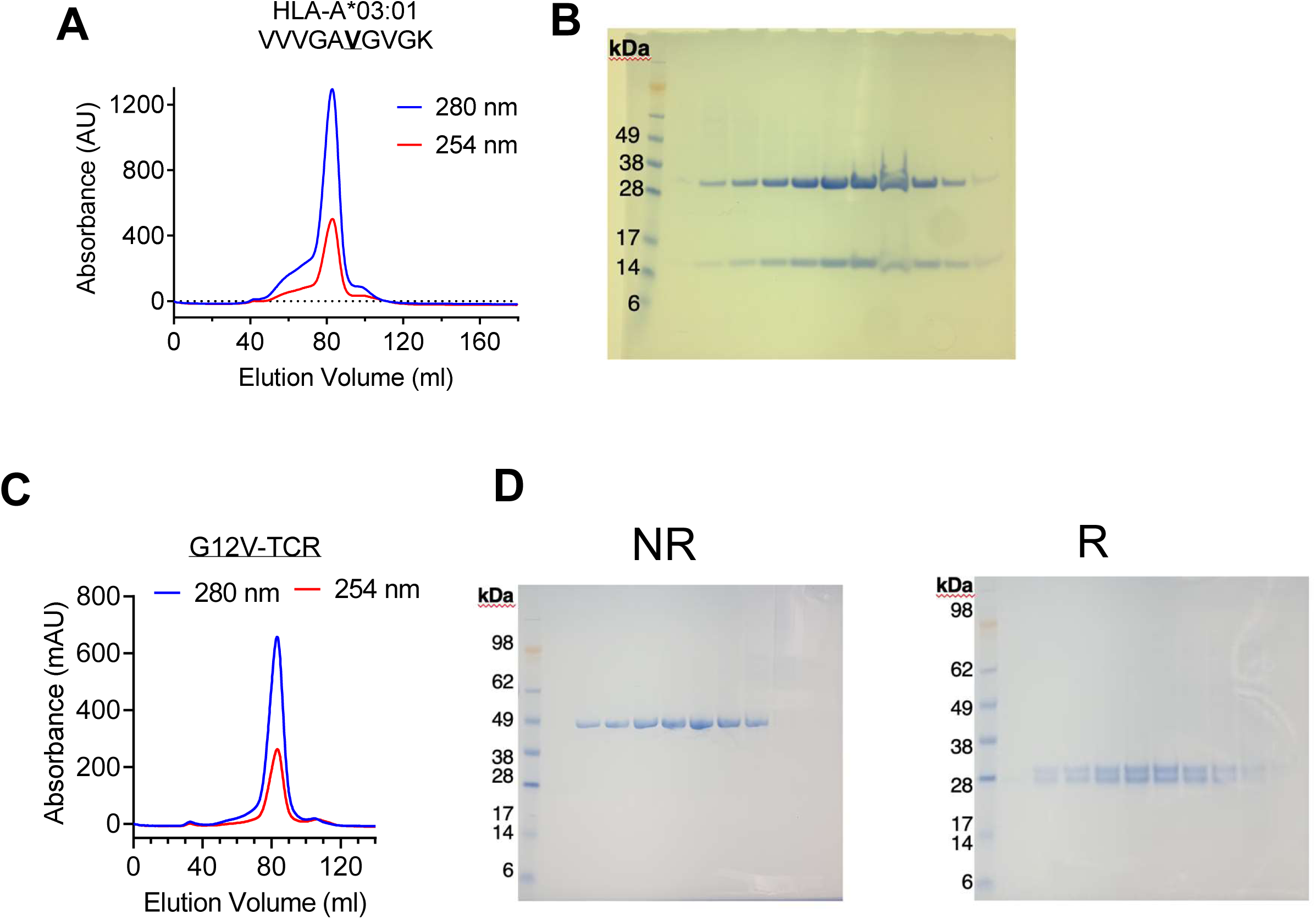
Production of recombinant HLA-A3 and G12V-TCR. (**A**) Representative size exclusion trace for recombinant HLA-A3 refolded with G12V-KRAS-10mer. Absorbance readings at 280nm and 254nm are shown. Column was Superdex S200. **(B)** Reducing SDS PAGE of peak fractions in A stained with Coomassie blue**. (C)** Representative size exclusion trace for recombinant refolded G12V-TCR. Absorbance readings at 280nm and 254nm are shown. Column was Superdex S200. **(D)** Reducing (right) and non-reducing (left) SDS PAGE of peak fractions in C stained with Coomassie blue.

**SFigure. 2.**
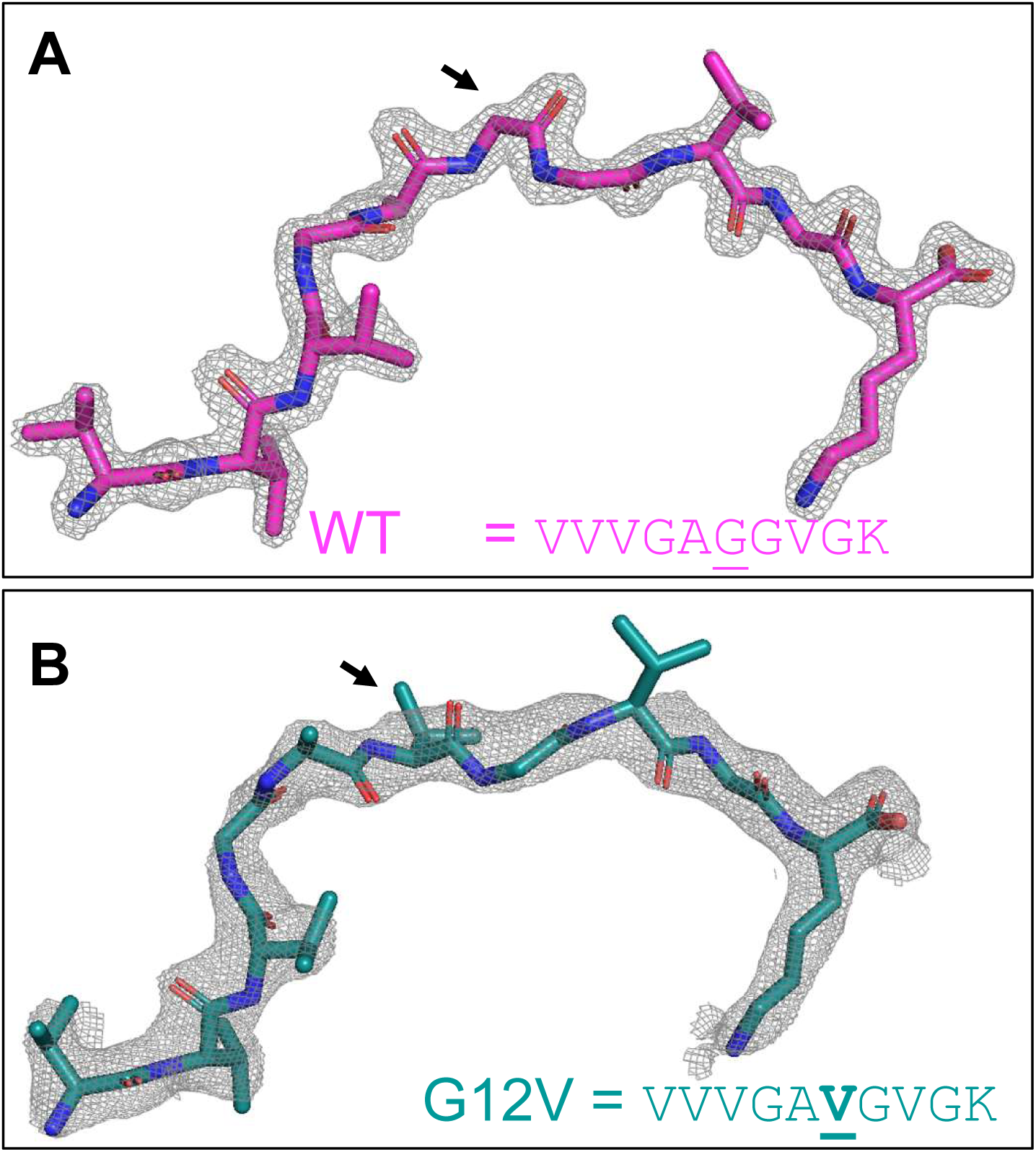
Structures of WT-KRAS and G12V-KRAS peptides. 2Fo-Fc omit map of WT-KRAS-10mer **(A)** and G12V-KRAS-10mer **(B)** contoured to 2s and 1.5s, respectively. Position 6, site of G12V mutation is indicated. Image generated in Pymol. PDB codes: 8VJZ (WT) and 8RNI (G12V).

**SFigure. 3.**
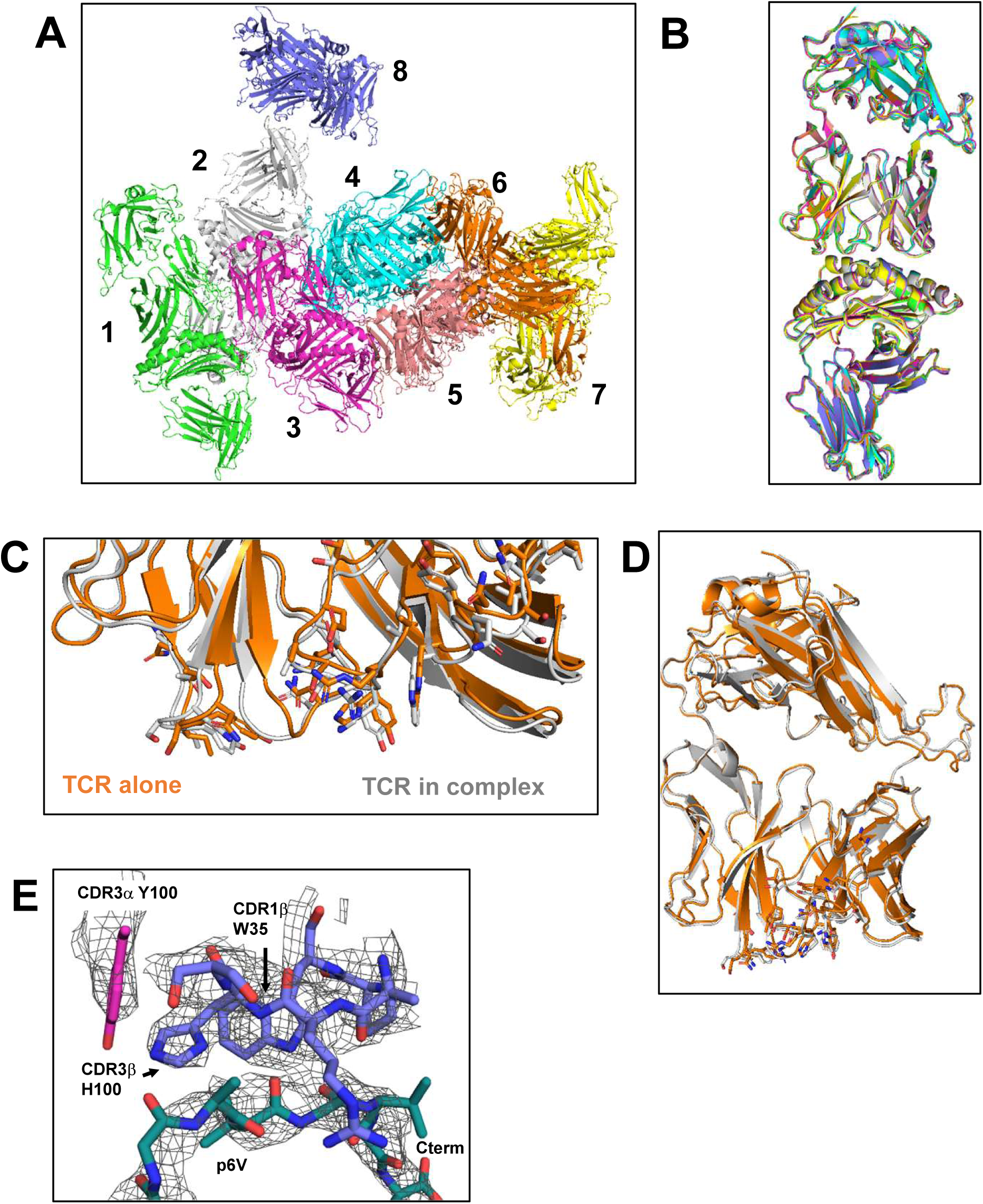
Structures of G12V-TCR alone and in complex with HLA-A3(G12V). (**A**) All 8 TCR-HLA-A3(G12V) complexes in the asymmetric unit. **(B)** All other 7 TCR-HLA-A3(G12V) complexes aligned with complex (chain ids: efghi). **(C&D)** Alignment of G12V-TCR structures determined alone (orange, PDB: 8RO5) and in complex with A3-G12V (grey, PDB: 8RRO). **(E)** 2Fo-Fc omit map of TCR-HLA-A3(G12V) complex displaying the G12V-10mer peptide, and major TCR contacts. Omit map is contoured to 1.5s.

**SFigure. 4.**
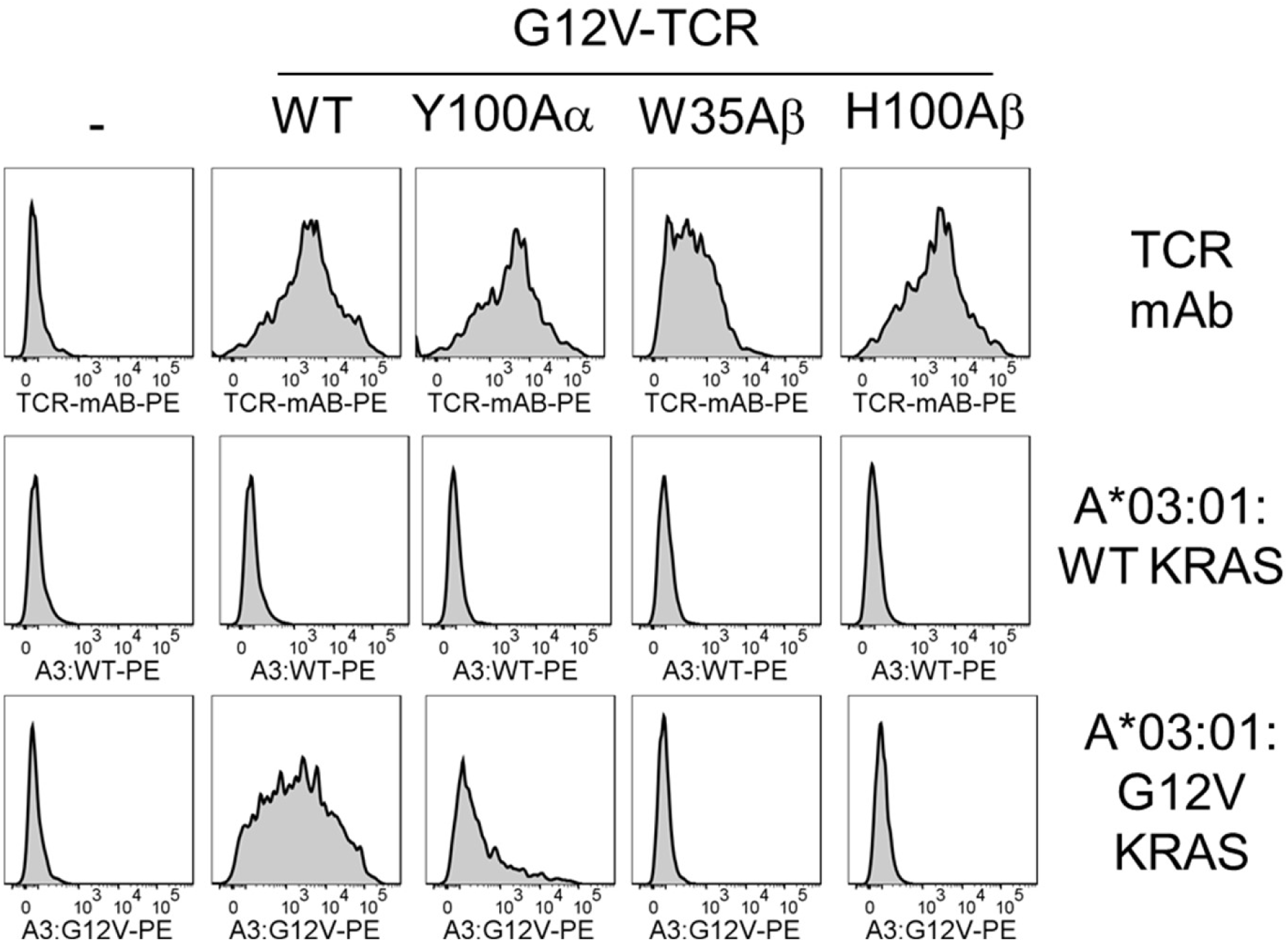
TCR expression in and HLA-A3 tetramer binding to G12V TCR transfected cells. 293T-CD3 cells were transfected with unmodified G12V-TCR and substitutions CDR3a Y100A, CDR1b W35A and CDR3b H100A. Cells were stained with TCR mAb, and HLA-A3 tetramers loaded with WT-KRAS (VVVGAGGVGK) and G12V-KRAS (VVVGAVGVGK) 10mer peptides.

**SFigure. 5.**
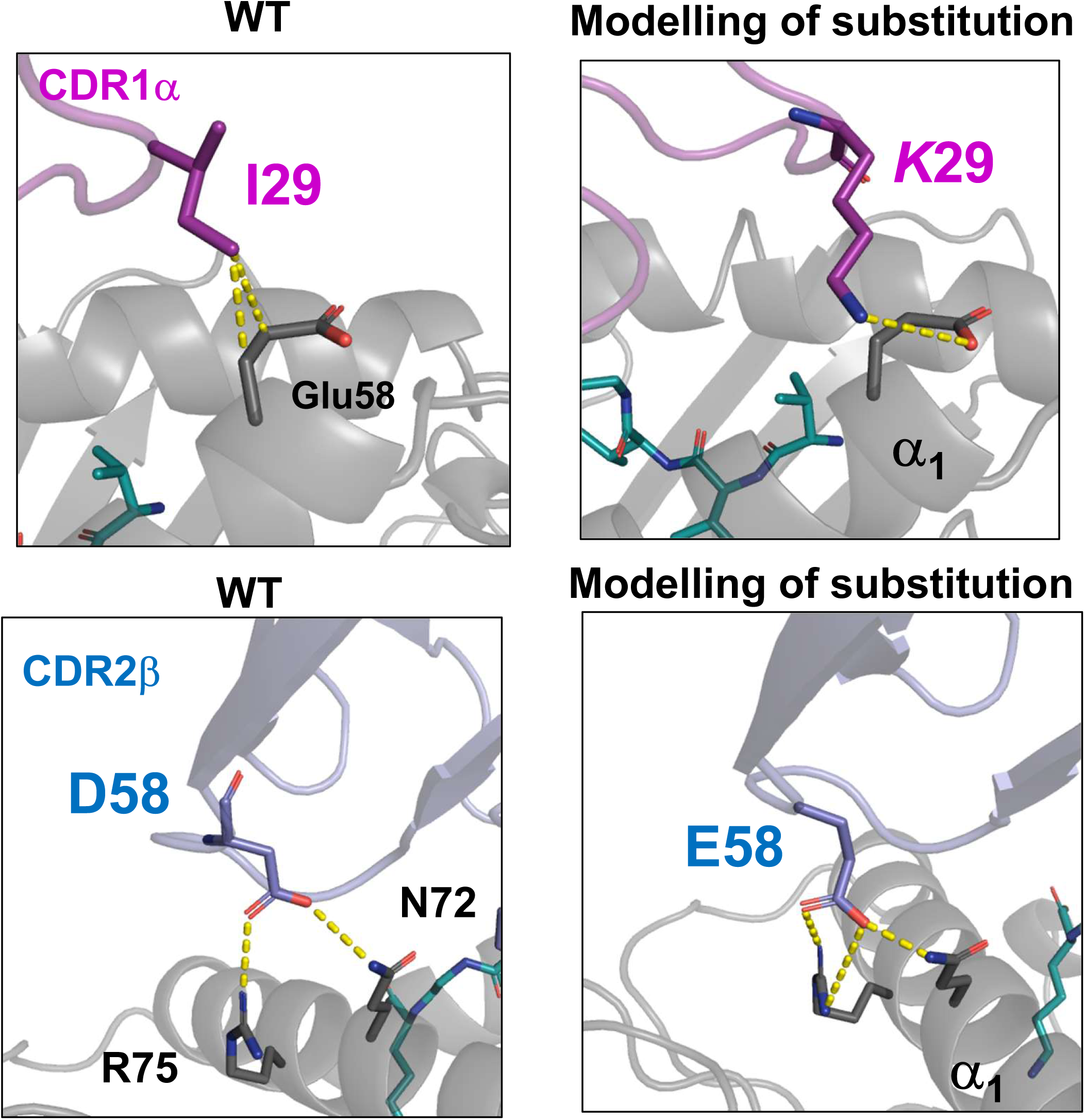
Modelling of TCR substitutions to increase binding strength. TCR contacts with HLA-A3 a1 helix (*left*) predicted to be improved with modelled substitutions (*right)*. Substitutions I29K (*top*) in CDR1a and D58E (*bottom*) in CDR2b are shown. Images and modelling were carried out in pymol.

**SFigure. 6.**
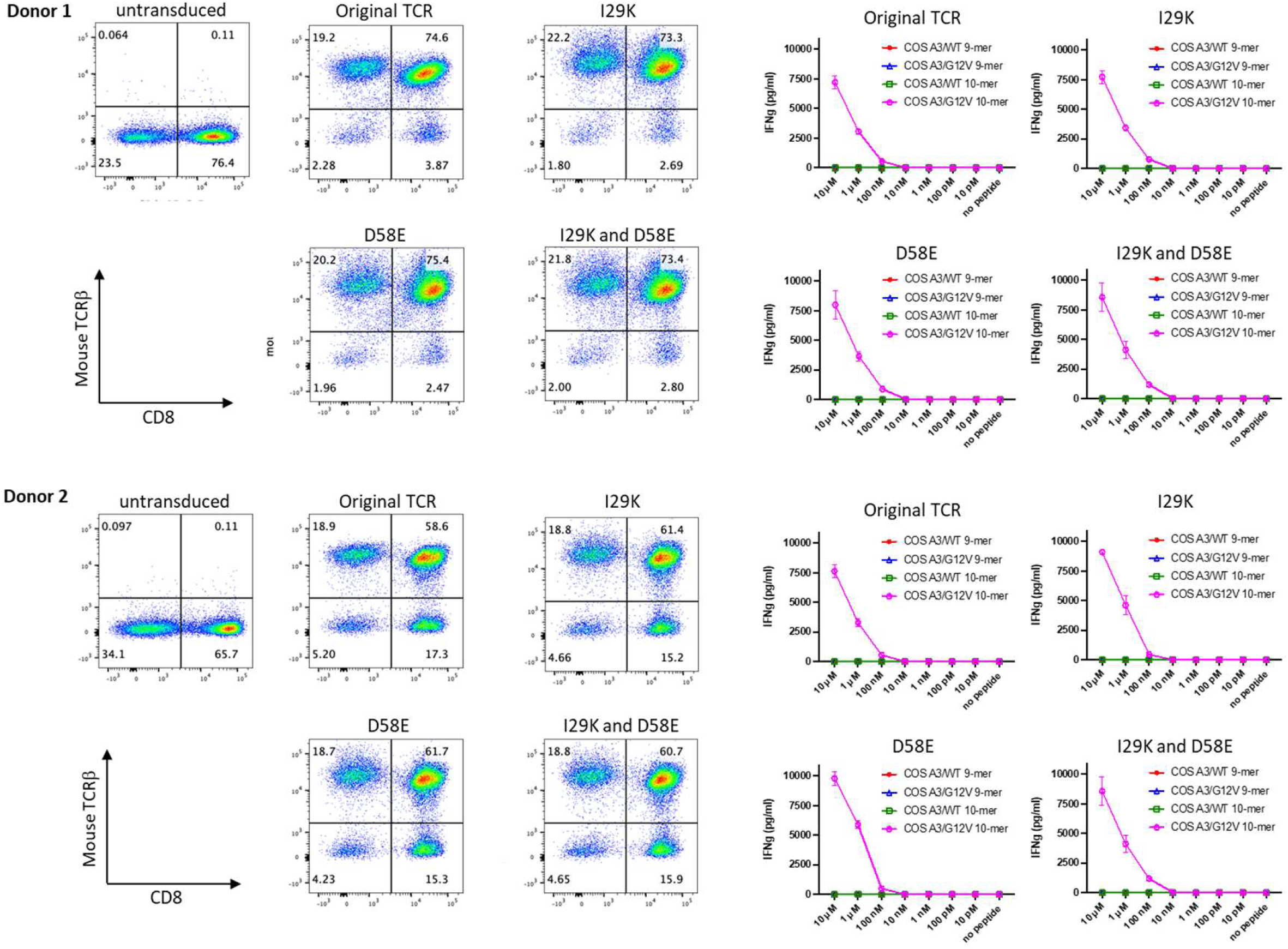
Functional test of modified TCRs. Genes encoding modified TCRs were cloned into retroviral vectors. These TCR genes were retrovirally introduced into peripheral blood T cells from two donors. TCR transduction efficiency was assessed by flow cytometry, and their sensitivity/specificity were tested by IFN-g ELISA using COS7/A3 cells pulsed with KRAS wild-type and G12V peptides at concentrations ranging from 10 µM to 10 pM as targets.

