## Supplemental Table 1 for "Identification and Structural Characterization of a mutant KRAS-G12V specific TCR restricted by HLA-A3"

| Structure | HLA-A3-WT-KRAS | HLA-A3-G12V-KRAS | G12V TCR | TCR-HLA-A3-G12V complex |
|---|---|---|---|---|
| **PDB code** | 8VJZ | 8RNI | 8RO5 | 8RRO |
| **Data collection** | | | | |
| Resolution range | 35.99 - 1.9 (1.968 - 1.9) | 42.67 - 2.49 (2.579 - 2.49) | 40.2 - 1.701 (1.762 - 1.701) | 46.31 - 3.5 (3.626 - 3.5) |
| Space group | P 6 2 2 | P 6 2 2 | P 1 21 1 | P 21 21 21 |
| *Cell dimensions* | | | | |
| a, b, c (Å) | 154.6 154.6 85.4 | 155.3 155.3 85.3 | 68.6 43.3 84.2 | 131.477 195.787 442.856 |
| $\alpha$, $\beta$, $\gamma$ (°) | 90 90 120 | 90 90 120 | 90 108.504 90 | 90 90 90 |
| Total reflections | 53078 (5023) | 24906 (2309) | 54445 (5244) | 147977 (11589) |
| Unique reflections | 47684 (4607) | 21743 (2052) | 51656 (5022) | 138025 (11066) |
| Multiplicity | 1.1 (1.1) | 1.1 (1.1) | 1.1 (1.0) | 1.1 (1.0) |
| Completeness (%) | 99.49 (98.86) | 98.83 (96.34) | 99.06 (97.36) | 84.39 (38.69) |
| Mean I/sigma(I) | 32.18 (7.71) | 20.76 (2.03) | 16.88 (3.10) | 3.46 (0.25) |
| Wilson B-factor | 22.65 | 43.76 | 17.63 | 110.3 |
| R-merge | 0.125 (0.653) | 0.23 (1.376) | 0.097 (0.506) | 1.408e-17 (2.539e-17) |
| R-meas | 0.127 (0.661) | 0.233 (1.408) | 0.104 (0.547) | 1.991e-17 (3.591e-17) |
| R-pim | 0.020 (0.106) | 0.036 (0.292) | 0.038 (0.207) | 1.408e-17 (2.539e-17) |
| CC1/2 | 0.989 (0.976) | 0.998 (0.764) | 0.993 (0.889) | 1 (1) |
| CC* | 0.997 (0.994) | 1.00 (0.391) | 0.998 (0.970) | 1 (1) |
| **Refinement** | | | | |
| Reflections used in refinement | 47484 (4607) | 21532 (2053) | 51502 (5022) | 137380 (5536) |
| Reflections used for R-free | 2430 (218) | 1070 (100) | 2657 (261) | 6098 (264) |
| R-work | 0.1873 (0.2349) | 0.2268 (0.3603) | 0.1830 (0.2563) | 0.2372 (0.3817) |
| R-free | 0.2235 (0.2801) | 0.2763 (0.4250) | 0.2094 (0.3099) | 0.2864 (0.4002) |
| CC(work) | 0.957 (0.925) | 0.952 (0.617) | 0.959 (0.909) | 0.945 (0.447) |
| CC(free) | 0.948 (0.885) | 0.920 (0.416) | 0.948 (0.770) | 0.910 (0.460) |
| Number of non-hydrogen atoms | 3575 | 3222 | 3903 | 53217 |
| macromolecules | 3163 | 3137 | 3552 | 53216 |
| ligands | 18 | 36 | 0 | 0 |
| solvent | 394 | 49 | 351 | 1 |
| Protein residues | 386 | 386 | 444 | 6595 |
| RMS(bonds) | 0.007 | 0.009 | 0.007 | 0.012 |
| RMS(angles) | 0.92 | 1.01 | 1 | 1.53 |
| Ramachandran favored (%) | 98.68 | 95.26 | 98.41 | 96.24 |
| Ramachandran allowed (%) | 1.32 | 4.74 | 1.59 | 3.76 |
| Ramachandran outliers (%) | 0 | 0 | 0 | 0 |
| Rotamer outliers (%) | 0.3 | 2.4 | 0.51 | 0.38 |
| Clashscore | 3.39 | 8.25 | 3.3 | 14.61 |
| Average B-factor | 28.75 | 48.99 | 21.5 | 123.4 |
| macromolecules | 27.92 | 49.03 | 20.96 | 123.4 |
| ligands | 41.48 | 56.9 | | |
| solvent | 34.85 | 41.09 | 26.94 | 30 |
